# Exploring Essential Oils and Their Bioactive Components from Mediterranean Medicinal Plants as Natural Antigiardial Agents

**DOI:** 10.1101/2025.10.07.680899

**Authors:** Sara Marcos-Herraiz, Sara Alonso Fernández, María José Irisarri, Jaime Arroyo Díaz, Francisco Ponce-Gordo, Azucena González-Coloma, Juliana Navarro Rocha, Iris Azami-Conesa, María Teresa Gómez-Muñoz, María Bailén

## Abstract

Lamiaceae and Asteraceae plant species have been widely used in traditional medicine to treat gastrointestinal disorders in countries of the Mediterranean basin. Other properties, such as antioxidant, anti-inflammatory, anti-bacterial, anti-parasite, and anti-virus have been described. *Giardia duodenalis* is the most frequent intestinal protozoon affecting young children worldwide and nitroimidazoles are the main treatment, but non-responding cases are widely reported. In this paper we tested the antigiardial activity of several Lamiaceae and Asteraceae plants.

Essential oils from 22 traditional medicinal plants from Lamiaceae and Asteraceae families and its main components were tested against the parasite employing the MTT method. The composition of the essential oils was determined by a metabolomic approach (GC-MS). Transmission Electron Microscopy was used to evaluate morphological alterations.

The best antigiardial activity was obtained with the genera *Lavandula*, *Mentha*, *Thymus,* and *Satureja*, while the best selective indexes were obtained with compounds γ-terpinene, caryophyllene oxide, carvacrol and thymol. Synergies were observed with linalyl acetate + linalool (*Lavandula* EOs), linalyl acetate + ρ-cymene or thymol, or combinations of ρ-cymene, γ-terpinene, thymol, and carvacrol (Satureja EOs). Membranolysis, enlarged periplasmic vacuoles, and loss of the cytoplasmic content were evident effects of γ-terpinene on trophozoites after 1 hour.

The present study provides phytotherapeutic evidence of essential oils from species of *Lavandula*, *Mentha*, *Thymus,* and *Satureja* as natural antigiardial agents, as well as some of their main components, γ-terpinene, caryophyllene oxide, carvacrol and thymol.

## 1. Introduction

*Giardia duodenalis* (*G. duodenalis*) is the most frequent parasite of the gastrointestinal tract of humans, particularly children between 0 and 2 years of age, ranging 37.7% to 96.4% of prevalence depending on the location, and more than 40% of repeated infections (Rogawski et al., 2017). It is more prevalent in developing countries, but in the EU more than 18,000 cases of giardiasis were diagnosed in 2019 (ECDC, 2022). The infection can be asymptomatic, especially in the case of adult and immunocompetent individuals, but it is frequently accompanied by several clinical signs, such as severe abdominal pain, diarrhea, vomiting, malabsorption, anorexia, flatulence, weight loss, and some patients develop postinfectious irritable bowel disease and chronic fatigue (Mørch and Hanevik, 2020). In summary, an estimated 184 million symptomatic cases per year occur worldwide (Havelaar et al., 2015).

Treatment is based on nitroimidazoles, but during the last decades a grow in the number of refractory giardiasis (giardiasis that do not respond to treatment) has been reported. Some of the refractory cases are related to deficient immune responses, but there are also cases considered as resistant to the employed treatment (Nabarro et al 2015, Carter et al., 2018). This fact is relevant to returning travelers, in whom treatment resistances were reported as high as 50% of positive cases, especially in travelers returning from Asia (Nabarro et al., 2015). Besides, there is robust evidence of cross-resistance among similar compounds, i.e., 5-nitroimidazoles (Upcroft & Upcroft, 2001). In addition, a complication of some treated cases is the development of inflammation due to the dysbiosis provoked by metronidazole (Pilla et al., 2020). With this scenario, it is essential to find alternatives or complementary treatments for giardiasis.

Natural products are composed of secondary metabolites that help living organisms to defend against pathogens and environmental stress. Products extracted from plants have been employed successfully for centuries against disease, and many of the components displayed anti-inflammatory, antioxidant, anti-bacteria, anti-virus and anti-parasite properties (Aziz et al 2018, Ramsey et al 2020). In traditional medicine, the use of Lamiaceae and Asteraceae natural products in the management of gastrointestinal disorders predominates over its therapeutic use in other medical conditions, especially in countries of the Mediterranean basin, such as Morocco and Pakistan (Jamila and Mostafa, 2014, Rahman et al., 2016, Es-Safi et al., 2020, Redouan et al., 2022). Among these disorders, diarrhea and virus, bacteria and parasite infections are frequently mentioned. Plant-based anti-*Giardia* (AG) products are very promising as alternative and complementary treatment resources (Alnomasy et al., 2021). Lamiaceae and Asteraceae plants were the most widely employed (30% and 13.5%, respectively), while Apiaceae, Myrtaceae, Amaryllidaceae, Cucurbitaceae and Zingiberaceae were less frequently explored (4.5-10.5% of the studies). In the search of AG agents, Lamiaceae are also interesting since it comprises plants that are active against more than one parasite, such as *Melissa officinalis*, *Origanum vulgare*, or *Thymus vulgaris* (Anthoy et al., 2005). Most of the studies performed with plants employed aerial parts, including leaves, but seeds, fruits, grain, stems, peel, flowers, bulbs and other minor components have been explored. Among the extracts, aqueous extract and essential oils (EOs) predominate in the studies (30% and 25.4% of the studies), although there are also other extraction methods, such as ethanolic, methanolic, hydroalcoholic, chloroform, petroleum-ether, and hexane extracts (Alnomasy et al., 2021).

Some of the most active natural products against *Giardia* trophozoites and less cytotoxic were extracted from the Lamiaceae family, such as the dichloromethane, methanolic and hexane extracts of *Mentha x piperita* with IC_50_ values 0.8-9 µg/ml (Vidal et al 2007), ethanolic extract of *Menta longifolia* and chloroform extract of *Ocimum basilicum* (IC_50_ = 68.2 µg/ml and 53.3 µg/ml, respectively) (El-Badry et al., 20210), or the EOs from *Thymus capitata Origanum virens*, and *Thymus zygis* subsp. *sylvestris*, with IC_50_ between 71 and 190 µg/ml (Machado et al 2010a and b).

Extracts from Asteraceae have demonstrated good AG activity, although cytotoxicity studies are necessary. Among them, we can highlight the ethanolic crude extracts and EOs from *Ageratum conyzoides*, with IC_50_ values ranging 35-96 µg/ml (Pintong et al., 2020). Natural products from other plants, such as the EO from the Myrtaceae *Syzygium aromaticum*, also displayed an interesting activity against *Giardia* trophozoites, with IC_50_ = 134 µg/ml (Machado et al., 2011). Garlic (*Allium sativum*) extracts are less effective, with IC_50_ = 300 µg/ml (Harris et al, 2000) or IC_50_ > 400 µg/ml (Argüello-García et al., 2018), although some compounds obtained from garlic were highly effective against *Giardia* trophozoites (Argüello-García et al., 2018, Harris et al., 2020).

There are several extracts with promising results against *Giardia* spp. but not all of them identified or analyzed the AG activity nor the main components of the extracts. Excluding compounds with cytotoxic effects or without cytotoxicity assays, there are only a few compounds pointed out as alternatives to fight *Giardia*, considering compounds with IC_50_ doses equal or lower than 100 µg/ml. Linearolactone is a diterpenoid compound isolated from *Salvia* species (i.e. *S. gesneriflora*) from Mexico, with AG activity (IC_50_ =28.2 µg/ml) (Calzada et al., 2015). Also, eugenol, a major component of *Syzygium aromaticum* EO, displayed IC_50_ value of 101 µg/ml (Machado et al., 2011).

To increase the number of active natural products or compounds against *Giardia* spp., we analyzed the AG activity of 30 EOs from Lamiaceae and Asteraceae medicinal plants obtained by two extraction methods, hydrodistillation (HD) and steam distillation (SD), to explore their utility as alternative or complementary treatments against giardiasis. We also tested cytotoxicity and AG activity of their main components alone or in combinations to investigate possible synergies. Finally, we explored the morphological alterations produced by the most active compound.

## 2. Methods and material

### 2.1. Plants and Essential Oils extraction

Essential Oils were obtained from plants belonging to the families Lamiaceae and Asteraceae. In total, 30 EOs from 22 different plants were used in the present study, all of them were obtained by HD, while eight of them were also extracted by SD (Table 1) (*Lavandula* x “*intermedia*” varieties Abrial and Grosso, *Lavandula luisieri*, *Lavandula angustifolia, Lavandula angustifolia var. mallete, Mentha suaveolens*, *Origanum virens*, *Satureja montana*, *Thymus vulgaris*, *Thymus zygis*).

**Table 1.**
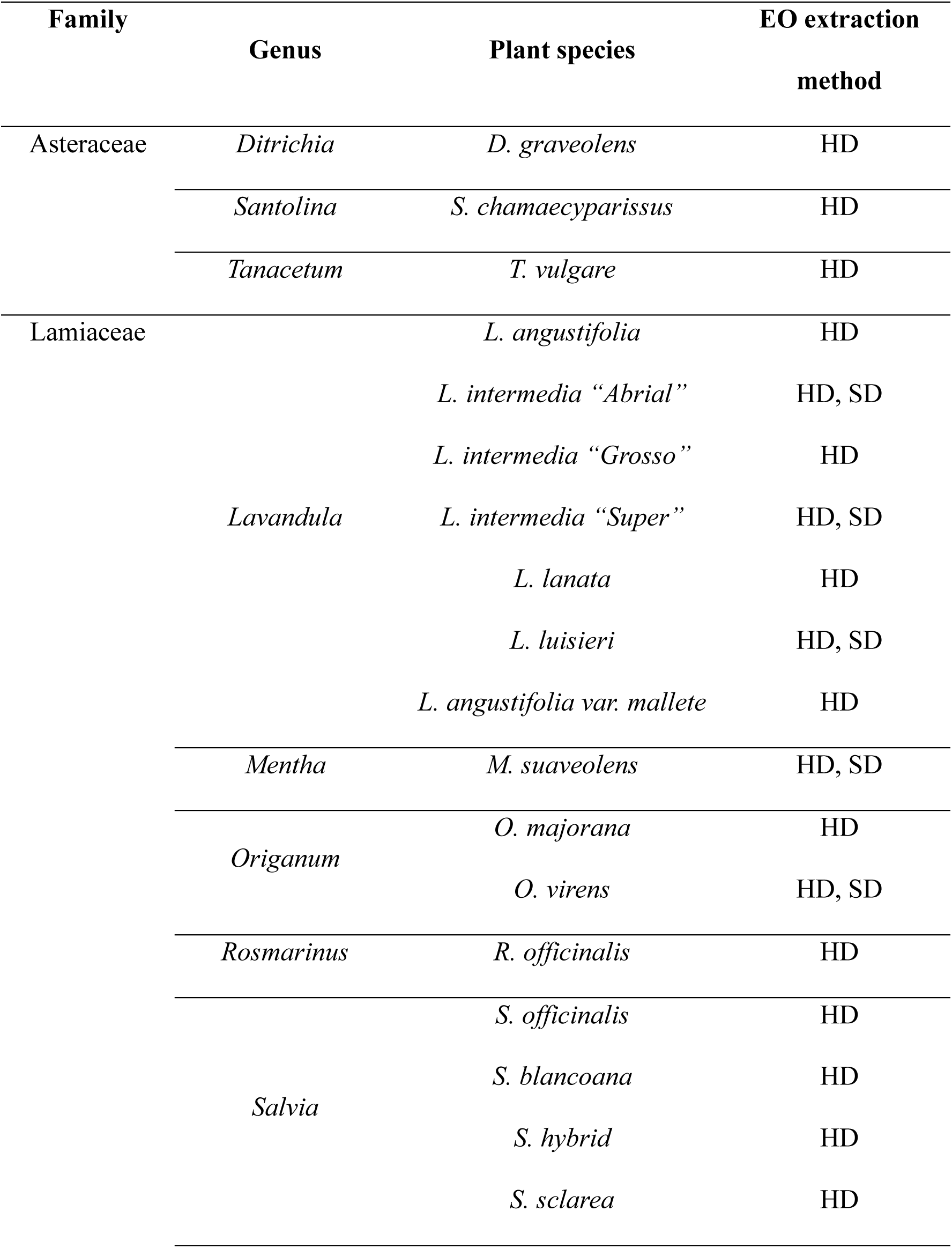
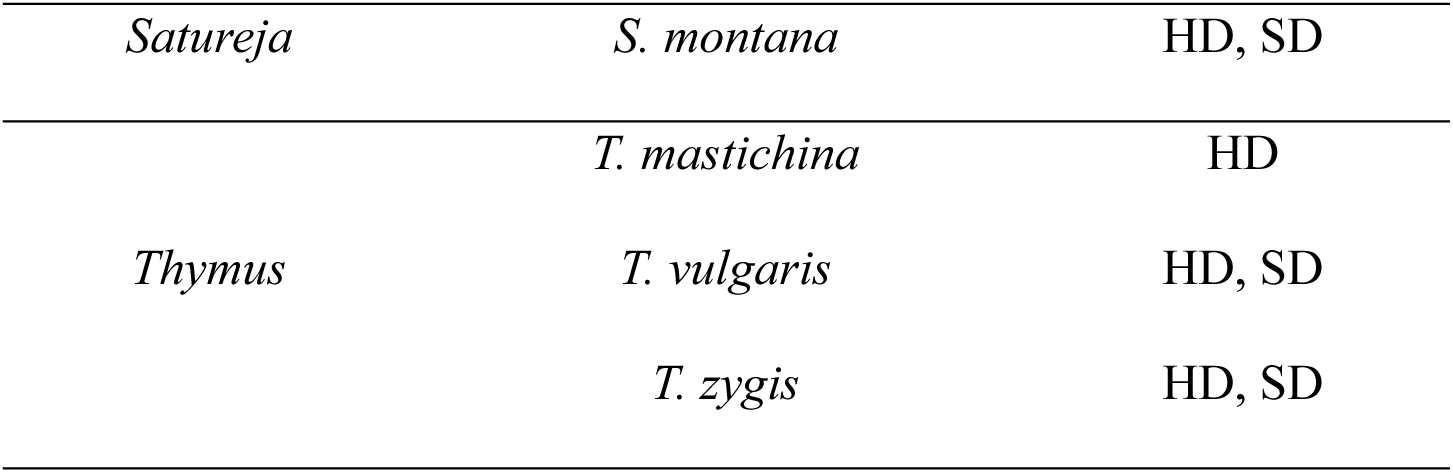
Families, plant species and extraction methods of the EOs employed in the present study.

All the plant species have been domesticated or are in culture. Seeds from the rest of the plants were obtained at the germplams banks of CITA (Centro de Investigación y Tecnología Agroalimentaria de Aragón, Plant Science Department, Zaragoza, Spain). The plant species were identified by Dr. Daniel Gómez (Instituto Pirenáico de Ecología, IPE-CSIC). Location, identification, and voucher numbers of the remaining species have been previously reported (Bailén et al., 2022). The plant names have been checked with http://www.theplantlist.org.

The aerial part of each plant was maintained during 7 days at room temperature and preserved from the light. The hydrodistillation process was conducted in triplicate employing 100 g of dried aerial part and 2 l of water in a Clevenger-type apparatus, according to the European Pharmacopeia recommended procedure (European Pharmacopeia). The oils were dried, filtered and stored at 4°C until use and determination of EOs composition. The steam distillation procedure was performed using 60 kg of fresh biomass harvested at the flowering stage. A stainless-steel pilot extraction plant with a reducing pressure valve was used, with a work pressure of 0.45 bar. The aqueous phase (hydrolate) was decanted from the EOs collected at the condensation section and filtered (Julio et al., 2017).

### 2.2. Identification of EOs main components

The main compounds of the EOs from the plants were identified by a metabolomic approach, using gas chromatography-mass spectrometry (GC-MS) employing a Shimadzu GC2010-Plus coupled to a Shimadzu GCMS-QP2010-Ultra mass detector with a single Quadrupole analyser. The electron impact ionization source was at 70 eV, employing Helium as gas carrier. The conditions have been described elsewhere (Bailén et al., 2022).

The mass spectra, retention time, and retention indexes were compared with data of the Wiley database (Wiley 275 Mass Spectra Database, 2001) and NIST 17 (NIST/EPA/NIH Mass Spectral Library). Quantification was conducted considering the relative area percentages of all peaks from the chromatograms. Trans-α-necrodyl acetate from *L. luisieri* was identified from a compound previously isolated.

### 2.3. Pure Compounds

After determining the composition of the analyzed EOs, compounds that were available and representing at least more than 5% of the active EOs, were employed in AG activity assays. Compounds not identified, not commercially available or not easy to isolate or to obtain were excluded.

Monoterpenes and sesquiterpenes were obtained from commercial sources, except for camphor that was isolated in the facilities of ICA-CSIC laboratory. Linalool, limonene, thymol, α-pinene, α-terpineol, β-pinene, camphene, β-caryophyllene and caryophyllene oxide were obtained from Sigma Aldrich (Madrid, Spain); linalyl acetate, γ-terpinene, and ρ-cymene from Acros Organics (Madrid, Spain); 4-terpineol from Merck Life Sciences (Madrid, Spain); α and β thujone from Phytolab; borneol, carvacrol, 1.8-cineole, D-fenchone, and ocimene from Fluka (Madrid, Spain). Metronidazole was purchased from Sigma Aldrich (Madrid, Spain).

### 2.4. Giardia duodenalis culture

Strain WB from *G. duodenalis* (obtained from a duodenal aspirate of a human) was purchased several years ago at the American Type Culture Collection (ATCC, isolate #30957™). The trophozoites were grown in Keysteŕs modified TYI-S-33 medium at 37°C in anaerobiosis. Trophozoites were grown in 15 ml polypropylene tubes, and the medium was refreshed every 2-3 days. When a monolayer was observed, the tube was cooled down at 4° C during 30-40 minutes for detachment and split in half. Trophozoites were preserved in 2 ml cryovials containing 5% dimethyl sulfoxide (DMSO) in culture medium at −80°C for medium term storage, or in liquid nitrogen for long-term storage.

### 2.5. Anti- *G. duodenalis* activity assays of EOs and pure compounds

The metabolic activity of the trophozoites was determined employing the MTT (3-(4,5-Dimethylthiazol-2-yl)-2,5-Diphenyltetrazolium Bromide) method (Beneré et al., 2007) with modifications (Arroyo Díaz et al 2024). Briefly, 145 µl of a solution containing 1×10^6^ trophozoites/ml in TYI-S-33 medium were seed in 96 well microtiter plates (Falcon BD, Fischer scientific, Madrid, Spain), and 5 µl of each EO or pure compound solution in DMSO (≤1%) were added in each well at different concentrations, in quadruplicate. EOs were evaluated at 800, 400, and 200 µg/ml, while pure compounds were tested at 100, 10 or 1 µg/ml. In case of significant AG activity of the EOs at 200 µg/ml, concentrations of 100, 50 and 25µg/ml were additionally evaluated. In the case of active pure compounds at least four intermediate concentrations between 100 and 1 µg/ml were also assayed to determine the inhibitory concentration of 50% (IC_50_) dose. Assays were incubated at 37°C during 24 hours, and then TYI-S-33 media was aspirated from each well and replaced for 100 µl of MTT-PMS (phenazine methosulfate, SIGMA, Madrid, Spain) solution (0.25 mg/ml MTT + 0.06 mg/ml PMS in PBS). After 75 min. At 37°C, 100 µl of sodium dodecyl sulphate were added to each well and incubated for 30 more minutes. After that, the plates were read in a spectrophotometer at 570 nm (Biotek 800ST microplate reader).

The AG activity was calculated as a percentage of the metabolic activity of the trophozoites, employing the formula: %AG=100-[(As-Ab)/(Ac-AB)]x100, where As is the absorbance of the sample (tested EO or compound), Ab is the absorbance of the blank (only culture medium), and Ac is the absorbance of the control (trophozoites culture without treatment). Metronidazol (Sigma Aldrich (Madrid, Spain) was employing as the reference drug for *Giardia*.

Those EOs with IC_50_ values lower than 200 µg/ml, according to previous criteria of promising AG activity (Machado et al., 2010b, Calzada et al 2015), were subjected to further analysis, while EOs with IC_50_ values higher than 200 µg/ml were discarded and they were not employed in more experiments.

### 2.6. Determination of synergisms in compounds combination

The synergistic activity of pure compounds was determined employing combinations of pure compounds from the most active EOs, following the method previously described, and employing a final concentration of 100 µg/ml (both compounds). The following combinations of paired products were tested at 1:1 (50µg/ml +50µg/ml), 2:1 (66.6µg/ml + 33.3µg/ml) and 1:2 (33.3 µg/ml + 66.6 µg/ml) proportions: linalyl acetate + 1,8-cineol; linalyl acetate + camphor; linalyl acetate + γ-terpinene; linalyl acetate + α-terpineol; linalyl acetate + ρ-cymene; linalyl acetate + thymol; linalyl acetate + linalool; linalool + 1,8-cineol; linalool + camphor; linalool + α-terpineol; thymol + γ-terpinene; thymol + ρ-cymene; thymol + carvacrol; ρ-cymene + γ-terpinene; ρ-cymene +carvacrol; γ-terpinene + carvacrol; camphor + 1,8-cineol. Non-available pure compounds, compounds with IC_50_ higher than 100µg/ml or compounds with cytotoxic activity were excluded from the synergism study.

Synergism was considered when the AG activity of the pair compounds assays (observed AG activity) was higher than the sum of the AG of each compound separately (expected AG activity) (Guardo et al., 2017).

### 2.7. Cytotoxicity assays and Selectivity Index

Cytotoxicity assays were conducted in 96 wells plates following the MTT method and employing Vero cells (a cell line derived from the kidney of African green monkey) (González-Coloma et al., 2002). Briefly, cells were cultured in 10% foetal calf serum (FCS, Sigma, Madrid, Spain) Dulbecco’s modified Eagle’s minimal essential medium (DMEM, Avantor, Llinars del Vallès, Barcelona) at 37°C and 5% CO_2_ in humidified atmosphere. Serial dilutions of the pure compounds were added to each well containing 1×10^4^ cells/well in quadruplicate. To determine the concentration that kill 50% of the cells, cytotoxicity concentration 50 (CC_50_), the compounds were employed at final concentrations of 100, 75, 50, 25, 10 and 1µg/ml.

The minimum selectivity index (SI) of the active compounds was calculated following the formula SI=CC_50_/IC_50_, considering that the highest CC_50_ assayed was 100 µg/ml. Those products with SI higher than 1 were considered as potential AG compounds.

### 2.8. Transmission Electron Microscopy

*Giardia* trophozoites treated with the most active pure compound against the parasite were processed for Transmission Electron Microscopy (TEM) analysis to look for ultra morphological alterations. Trophozoites treated with the compound, as well as control trophozoites without treatment, were processed after 60 min and 24h exposure.

*Giardia* cultures were cooled down at 4°C for 40 minutes to detach the trophozoites. The tubes were centrifuged at 800 x *g* for 5 minutes and washed with sterile cold PBS twice. The pellets containing the trophozoites were transferred to Eppendorf tubes for the fixation process. The concentrated parasites were fixed for 4 h at 4°C in Karnovsky buffer (4% paraformaldehyde, 2.5% glutaraldehyde Fisher scientific, Madrid, Spain) in Milloning phosphate buffer (Milloning, 1961). Then, trophozoites were washed four times with Milloning buffer, and left overnight at 4°C in 1 ml of the last washing buffer. After centrifugation, the trophozoites were post-fixed in 1% osmium tetroxide and potassium ferrocyanide 1.5% for 1 h, followed by three washing in Milli Q water. A dehydration procedure was conducted in 15 min steps of acetone in increasing concentrations (30%, 50%, 70%, 80%, 90%, 95%, 100%). The acetone was substituted by SPURR resin (Iberlabo S.A., Madrid, Spain) by washings in growing concentrations of resin: acetone 1:3 for 1h, 1:1 for 1 h, 3:1 for 2 h, and then pure resin overnight. The samples were embedded in new resin and were polymerized at 65°C for 48-72h. The embedded samples were sent to the Spanish National Centre for Electron Microscopy, where ultrathin sections (60 µm depth) where cut, loaded in a copper grid and examined in a JEOL JEM-1400 Electron Microscope operating at 80kV.

### 2.9. Statistical analysis

The data obtained was analyzed employing STATGRAPHICS Centurion 19. A linear regression analysis was employed to determine IC_50_ anti-*Giardia* activity, as well as CC_50_ cytotoxic activity (percentage of cell viability on log-dose).

The viability of cells and trophozoites was assessed for each compound through a dose-response experiment to determine their relative potency, expressed as CC_50_ or IC_50_ values. The IC_50_ (µg/mL) represents the concentration of EOs or pure compounds required to achieve 50% mortality of trophozoites, whereas the CC_50_ (µg/mL) indicates the concentration needed to induce 50% mortality in Vero cells.

The minimum selectivity index (SI) was calculated for pure compounds using the formula SI = IC_50_/CC_50_. Compounds with an SI greater than one were considered promising AG agents, as they exhibit higher toxicity toward protozoan than mammalian cells.

Normality of the variables was assessed using the Shapiro–Wilk test. Since the data were normally distributed, the Student’s t-test was applied.

## 3. Results

### 3.1. Anti-*Giardia* activity of EOs

The AG activity of the different EOs varies greatly with the plant species. All the EOs analyzed were active (80% AG activity) against *Giardia* at least at 800 µg/ml (Tables 2 and 3). Three plants extracted with both methods displayed similar IC_50_ values when HD and SD EOs were compared (*M. suaveolens*, *T. vulgaris* and *T. zygis*), three of them displayed better IC_50_ values with HD EO (*L. “Super”*, *L. “Abrial”* and *O. virens*), while only two plants displayed better results with the SD EO (*L. luisieri*, *S. montana*), although in the last case, both EOs good AG activity.

**Table 2.** Anti-*Giardia* activity of EOs from different plant species and extraction methods evaluated with IC_50_ values lower than 200 µg/ml (ordered from lower to higher IC_50_).

| Plant species | EO extraction method | IC<sub>50</sub> in µg/ml<br>(95% Confidence interval) |
| --- | --- | --- |
| *L. luisieri* | SD | <25 |
| *T. vulgaris* | SD | <25 |
| *T. vulgaris* | HD | <25 |
| *M. suaveolens* | SD | <25 |
| *M. suaveolens* | HD | <25 |
| *S. montana* | SD | <25 |
| *L. lanata* | HD | 27.9 (25.6-30.4) |
| *L. luisieri* | HD | 32.1 (24.2-42.7) |
| *T. zygis* | HD | 66.9 (65.1-68.7) |
| *T. zygis* | SD | 71.5 (62.4-81.9) |
| *S. sclarea* | HD | 88.2 (68.9-112.9) |
| *S. montana* | HD | 109.2 (100.9-118.2) |
| *Lavandula x intermedia* “Super” | HD | 111.3 (100.7-123.0) |
| *Lavandula x intermedia* “Abrial” | HD | 113.0 (100.9-126.4) |
| *O. virens* | HD | 199.4 (185.0-214.9) |

**Table 3.**
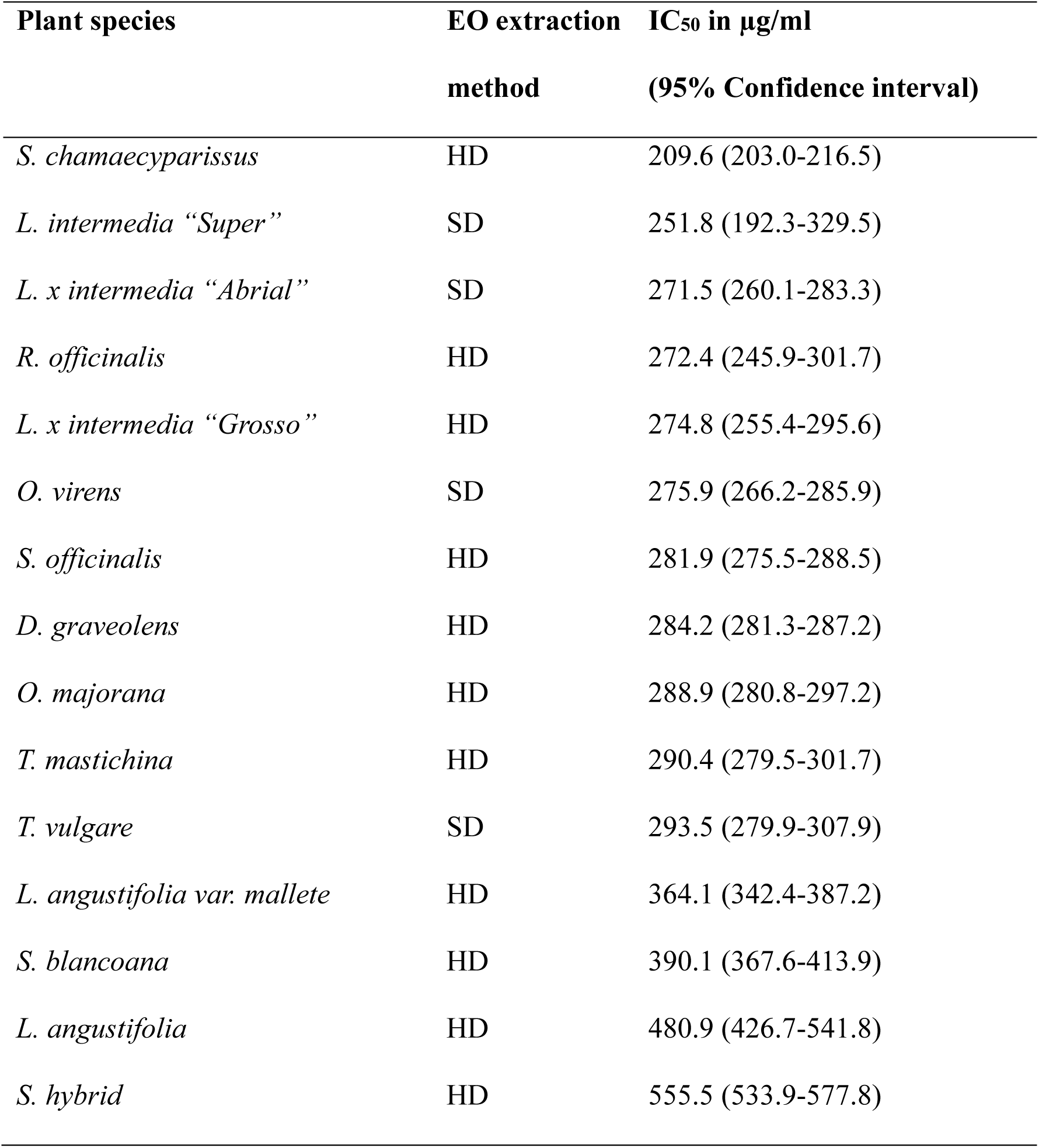
Anti-*Giardia* activity of EOs from different plant species and extraction methods evaluated with IC_50_ values higher than 200 µg/ml.

Considering the IC_50_ values, 15 EOs were selected for further analysis, since their IC_50_ values were lower than 200 µg/ml. They belonged to the genera *Lavandula*, *Thymus*, *Mentha*, *Satureja*, *Origanum* and *Salvia* (Table 2). The best IC_50_ values were obtained with *T. vulgaris* and *M. suaveolens* EOs obtained with both extraction methods, and *S. montana* and *L. luisieri* obtained with SD extraction method (IC_50_ value < 25 µg/ml).

The EOs with an IC_50_ higher than 200 µg/ml (n=15) were not subjected to further analysis (Table 3), except for the determination of their composition. IC_50_ values ranged from 209.6 to 555.5 µg/ml. These EOs were extracted from the following plants and methods of extraction: *L. angustifolia* HD, *L. intermedia* “Abrial” SD, *L. intermedia* “Grosso” HD, *L. intermedia* “Super” SD, *L. angustifolia var. mallete* HD, *D. graveolens* HD, *O. virens* SD, *O. majorana* HD, *R. officinalis* HD, *S. hybrid* HD, *S. officinalis* HD, *S. blancoana* HD, *S. chamaecyparissus* HD, *T. mastichina* HD, and *T. vulgare* SD.

### 3.2. Main components of the most active EOs

The main components of the different EOs will be explained from higher to lower activity against the parasite and comparing both extraction methods (Table 4, supplementary table 1). In general, the main components of the employed EOs were similar with both extraction methods, when employed.

**Table 4.**
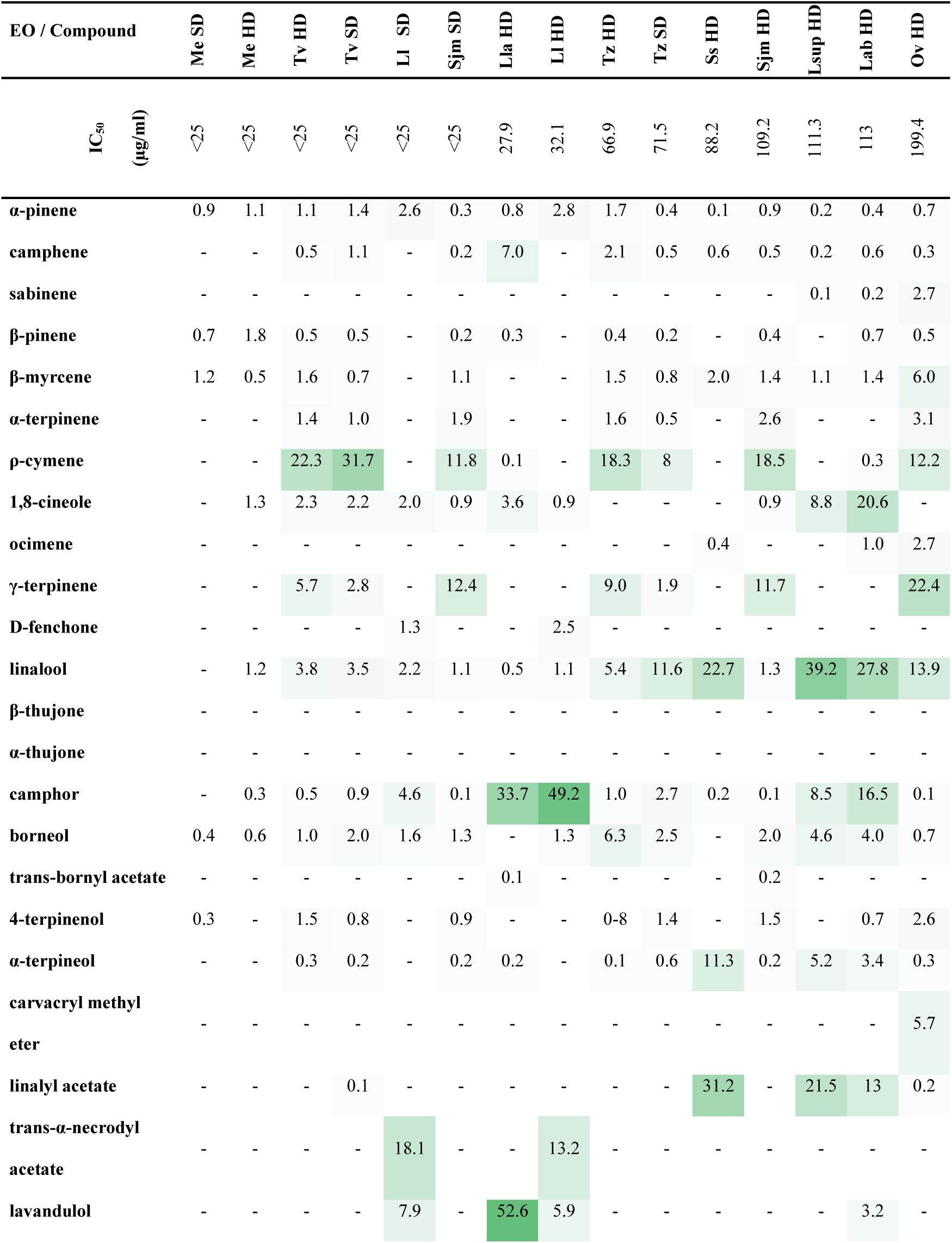
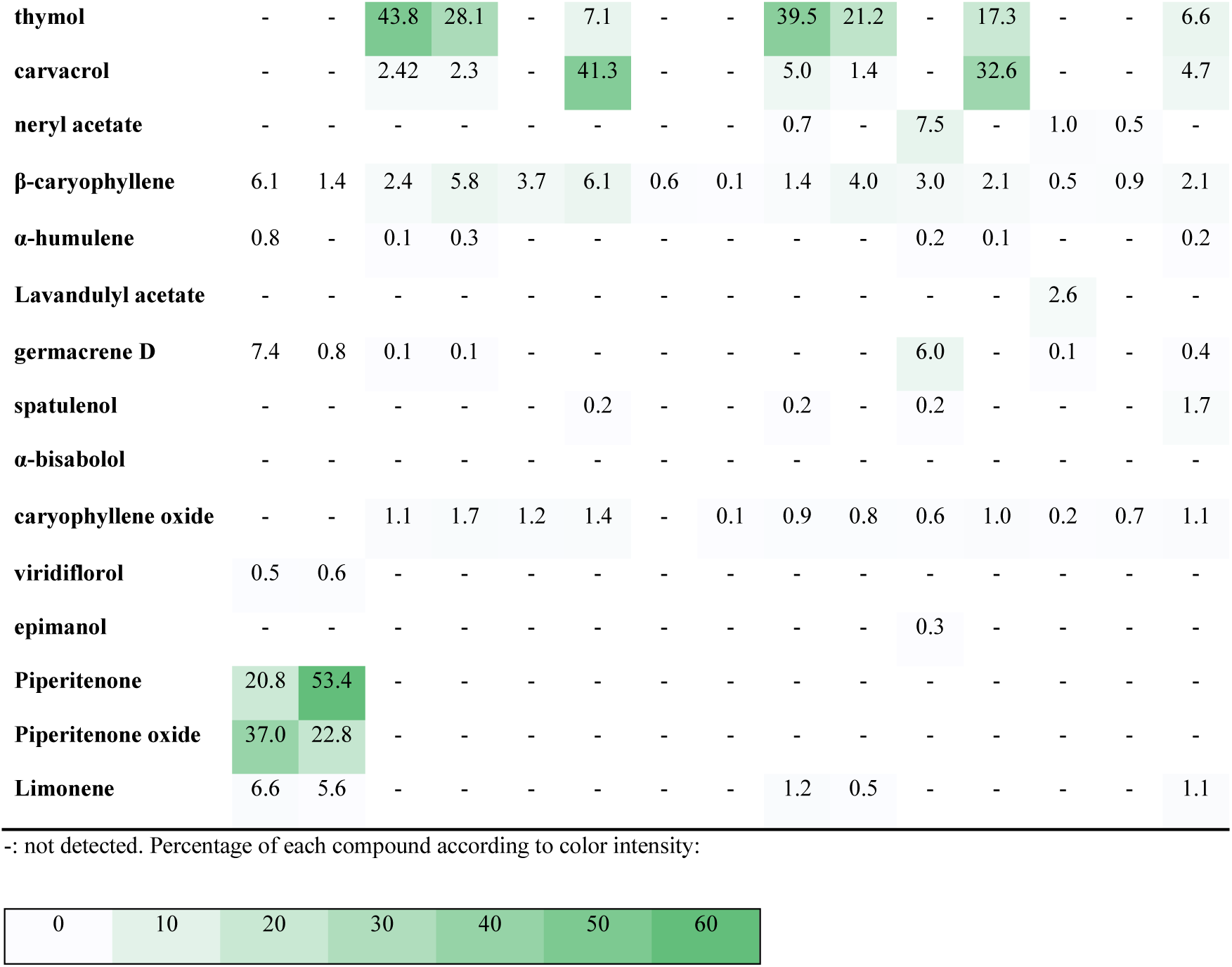
Main components (in percentage) of the EOs with IC_50_ values lower than 200 µg/ml and ordered by retention index. Me: *M. suaveolens*; Tv: *T. vulgaris*; Ll: *L. luisieri*; Sjm: *S. montana*; Lla: *L. lanata*; Tz: *T. zygis*; Ss: *S. sclarea*; Om: *O. majorana*; Lsup: *L. intermedia* “Super”; Lab: *L. intermedia* “Abrial”; Ov: *O. virens* HD, EOs obtained by hydrodistillation; SD, EOs obtained by steam distillation.

Piperitenone (20.7-53.4%) and piperitenone oxide (22.8-35%) were the most abundant components of *M. suaveolens* EOs, which displayed IC_50_ values lower than 25 µg/ml. At minor proportions, limonene (5.6-6.6%), germacrene D (0.8-7.4%) and β-caryophyllene (1.4-6.1%) were also identified (Table 3).

Two components predominate in the EOs from *L. luisieri*: camphor (4.6-49.5%) and trans-α-necrodyl acetate (12.4-18.8%). Lavandulol was present at low proportions (5.9-7.9%).

In the HD EO from *L. lanata*, with an IC_50_ of 27.9 µg/ml, lavandulol and camphor were identified at remarkably high proportions (52.6% and 33.7%, respectively).

The main components of the genus *Thymus* were similar in the four EOs extracted with two different methods. Thymol (21.2-43.8%) and ρ-cymene (8.0-31.7%) predominate in both plant species, although at higher concentrations in *T. vulgaris*, the most active of both plants (IC_50_ < 25 µg/ml). Other compounds present were γ-terpinene (1.9-9%) and linalool (3.5-11.6%) (Table 4).

Four components were identified at high proportions in the EOs of *S. montana* (IC_50_= <25-109.2 µg/ml), carvacrol being the most abundant (32.5-41.3%), followed by ρ-cymene (11.8-18.5), thymol (7.1-17.3%), and γ-terpinene (11.7-12.4%).

*Origanum majorana* HD EO (IC_50_= 288.9 µg/ml) revealed 4-terpinenol as the most abundant component (29.7%), followed by γ-terpinene (10.7%) and trans-caryophyllene, ocimene, ρ-cymene, α-terpinene, and sabinene with proportions between 5 and 6%.

The main components of *O. virens* HD EO, the most active (IC_50_ = 199.4 µg/ml), were γ-terpinene (22.4%), linalool (13.9%) and ρ-cymene (12.2%), while γ-terpinene (15.4%) and linalool (15.3%) predominate in the *O. virens* SD EO (IC_50_ value = 275.9 µg/ml). Minor components (4.7-6%) were also detected in *O. virens* HD EO (β-myrcene, carvacryl methyl ether, thymol, and carvacrol), with lower proportions in general in the SD EO.

In the HD extracted EO of *S. sclarea* (IC_50_ value = 88.2 µg/ml), a high amount of linalyl acetate (31.2%), linalool (22.7%), α-terpineol (11.3%), and neryl acetate (7.5%) were identified.

The EOs of *Lavandula x luisieri* were composed of four main components at different proportions: linalool (19.1-39.2%), linalyl acetate (13-37.8%), 1,8-cineole (7.9-20.6%), and camphor (6.7-16.5%). At minor proportions β-caryophyllene (5.5% in *L. intermedia* “abrial” SD EO) were also identified.

### 3.3. Anti-*Giardia* activity of main components, cytotoxicity and SI

Available pure compounds from the EOs were evaluated against *Giardia* trophozoites (n=16), most of them were present at concentrations higher than 5% in some of the active EOs, except three of them that were at concentrations higher than 2.5% (α-pinene, ocimene, 4-terpinenol). They were also assessed with Vero cells to analyze cytotoxicity (Table 5). Six compounds were active against *Giardia* trophozoites at doses lower than 100 µg/ml, α-pinene, γ-terpinene, caryophyllene oxide, β-caryophyllene, carvacrol and thymol, ordered from higher to lower AG activity (Figure 1). Of them, two showed cytotoxicity when evaluated at 100 µg/ml, α-pinene, β-caryophyllene. The other four compounds were considered as promising anti-*Giardia* agents, being γ-terpinene the compound with the highest SI obtained (2.4), followed by caryophyllene oxide (SI = 1.8), carvacrol (SI = 1.7), and thymol (SI = 1.3).

**Figure 1.**
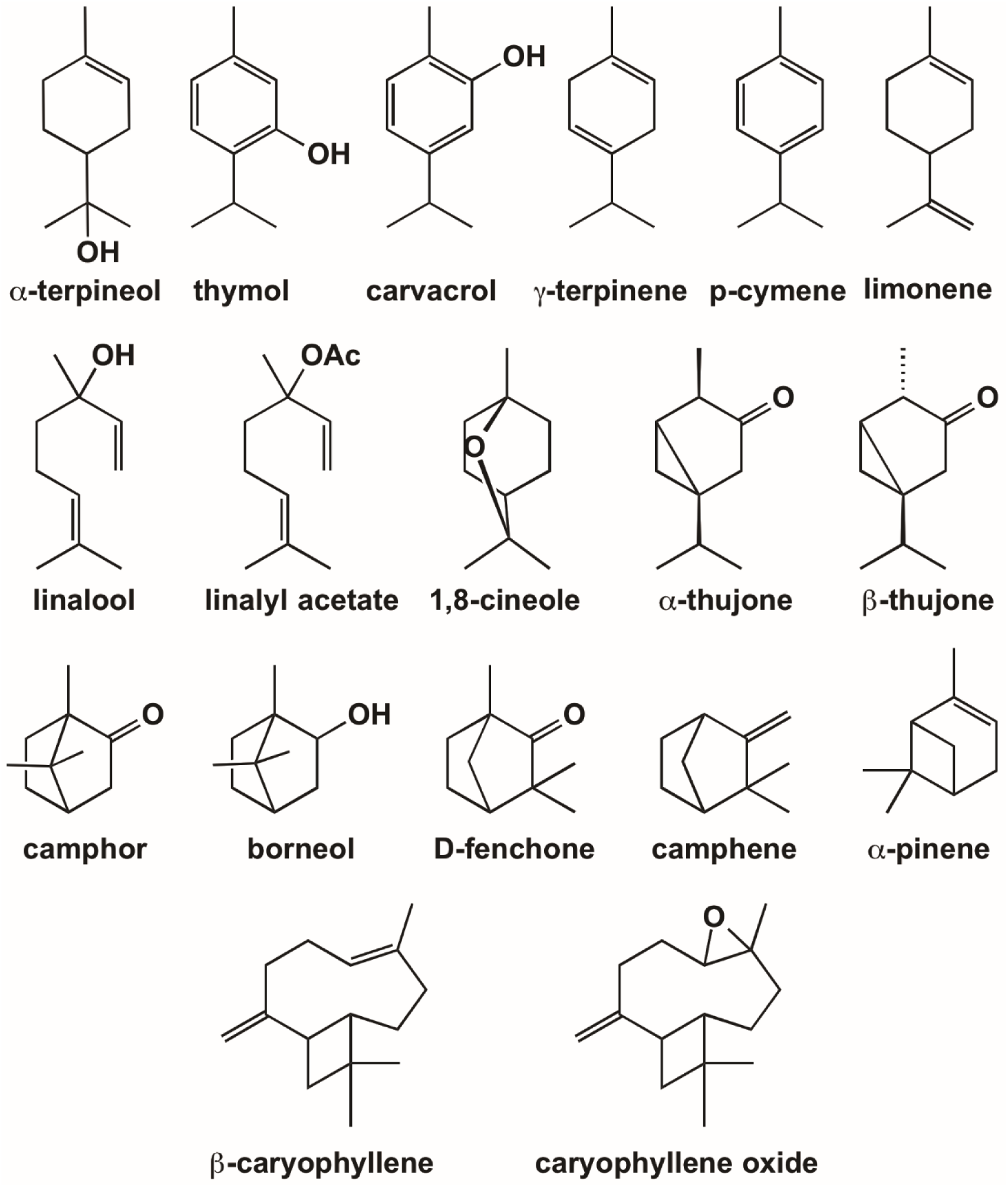
Chemical structure of the major components identified from the EOs employed in this study.

**Table 5.**
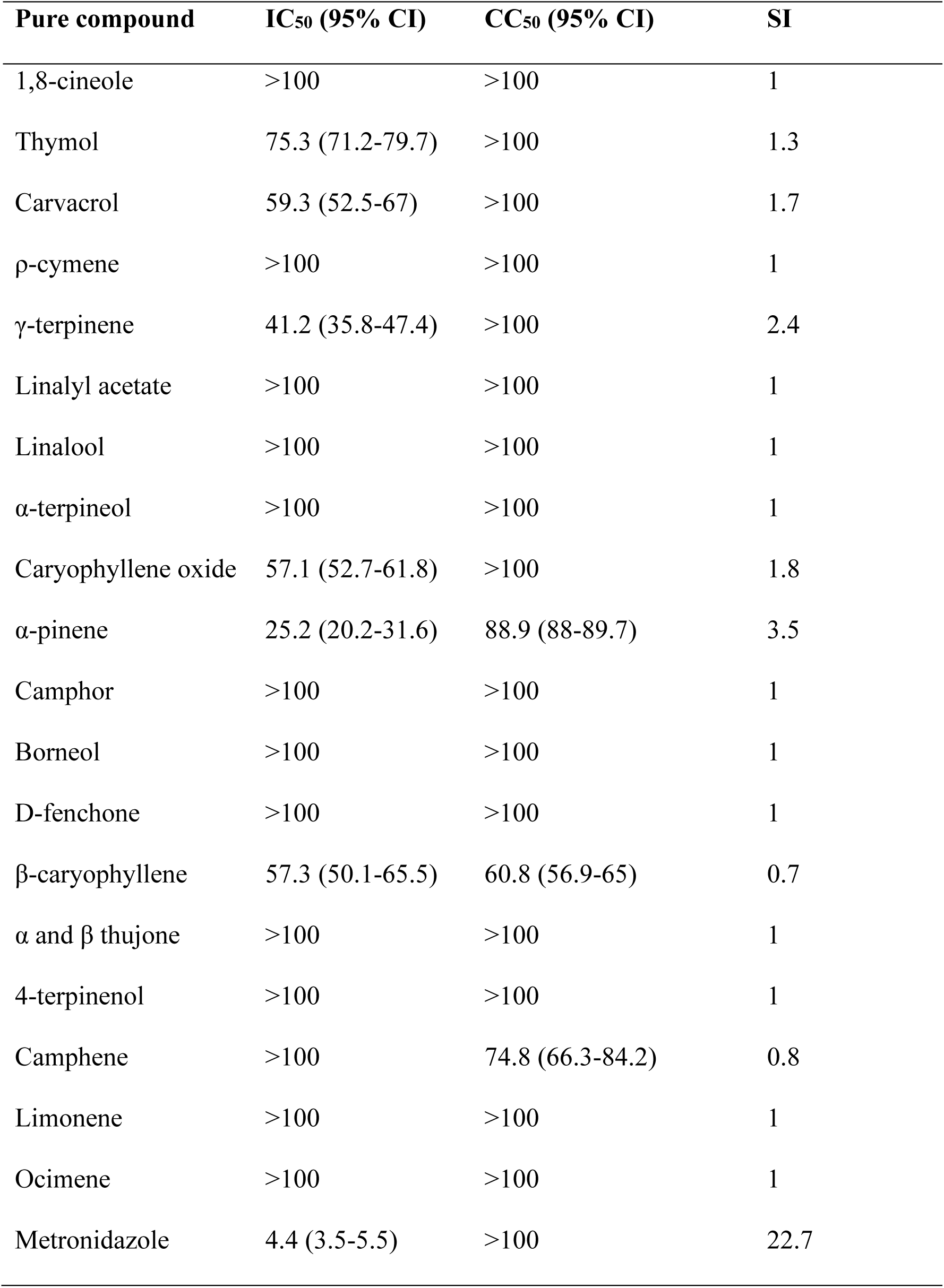
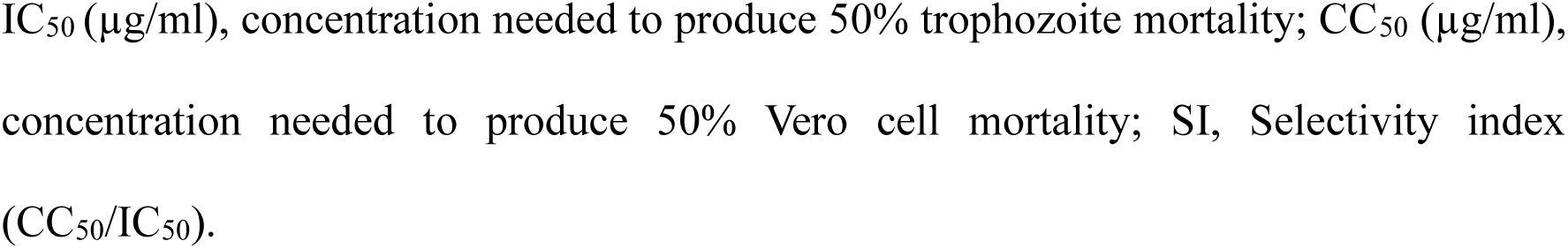
IC_50_ (µg/ml), CC_50_ (µg/ml) and SI values of the tested main components from the EOs.

### 3.4. Synergies of pure compounds

Considering the composition of the active EOs and the availability of pure compounds, their synergistic activity was explored by pair combinations of compounds (Figure 2, supplementary Figures 1 and 2). Compounds representing less than 8% of the composition and cytotoxic compounds were not considered for the synergies experiment.

**Figure 2.**
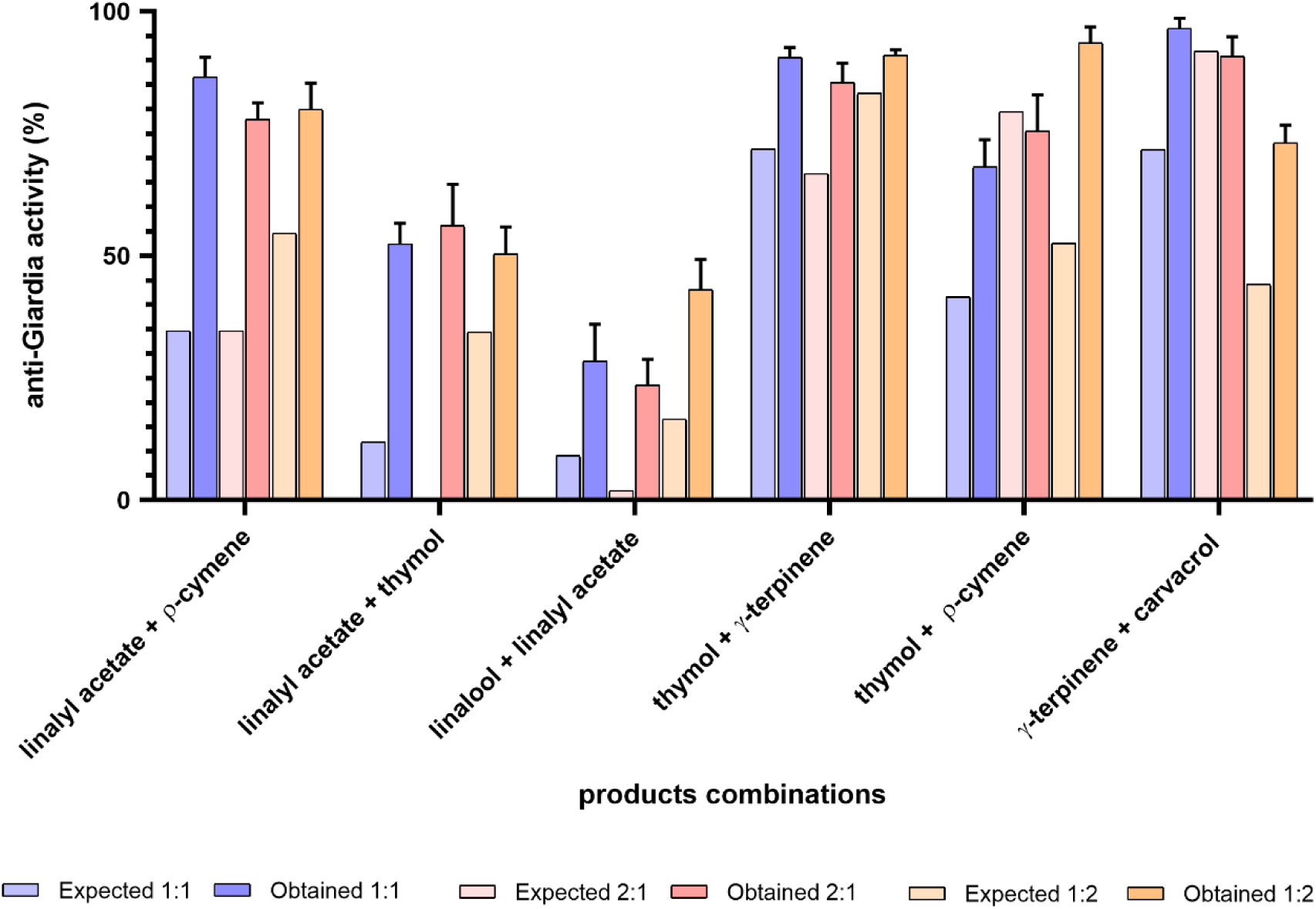
Activity of compounds combined in pairs from the most active AG EOs that showed synergistic activity.

The combinations of linalyl acetate with most of the AG active compounds (camphor, α-terpineol, 1,8-cineol, and γ-terpinene) did not show any synergistic activity. Also, combinations of linalool with most of the AG active compounds (camphor, α-terpineol, 1,8-cineol) did not show synergistic activity (Supplementary Figures 1 and 2).

However, the combinations linalyl acetate with the monoterpene linalool, as well as the combination linalyl acetate + thymol or ρ-cymene displayed synergistic activity since the observed AG activity was higher than the expected activity (sum of the activity of each compound separately) (Figure 2). The synergistic combinations linalyl acetate + linalool was present in several of the active EOs, *S. sclarea* HD EO (IC_50_ = 88.2 µg/ml), *L. intermedia* “super” HD EO (IC_50_ = 111.3 µg/ml), and *L. intermedia* “abrial” HD EO (IC_50_ = 113 µg/ml).

Synergistic effect was observed when thymol was combined with ρ-cymene or γ-terpinene, and when γ-terpinene was combined with carvacrol. These combinations were present in several of the most active AG EOs, such as *Thymus* and *Satureja* EOs (HD Tv, SD Tv, HD Tz, SD Tz, HD Sm, and SD Sm).

No synergistic effect was observed with the combination of carvacrol with thymol or ρ - cymene. These combinations were present in significant amounts in both *S. montana* EOs (IC_50_ of *S. montana* SD EO <25 µg/ml, and IC_50_ of *S. montana* HD EO = 109.2 µg/ml). However, besides a high amount of carvacrol, in both EOs γ-terpinene and thymol are present at amounts higher than 6%.

In the case of γ-terpinene, high activities (close to 100% both, in expected and obtained values) were observed when combined with ρ-cymene or linalyl acetate, and for that reason no synergistic activity could be determined (Supplementary Figure 2).

In summary, most of the synergies were observed when the monoterpenoid ester linalyl acetate was combined with monoterpenes or when monoterpenes were combined. The presence of thymol or ρ-cymene was observed in most of the cases of synergies, and carvacrol and γ-terpinene were observed in others. None of the other compounds evaluated were present in the synergies obtained.

### 3.5. Morphological changes of γ-terpinene on *G. duodenalis* trophozoites

Since γ-terpinene was the most active of the tested compounds against *Giardia* sp., morphological alterations of the trophozoites were evaluated employing TEM with the trophozoites non-exposed (Figure 3 A-C) or exposed (Figure 3 D-L) to the compound. Vacuolization and mortality were observed soon after exposition of γ-terpinene for 60 min, a moment at which most of the trophozoites seemed dead (Figure 3 D-F). Trophozoites displayed significant morphological alterations with respect to the non-exposed trophozoites, mainly consisting in broken plasmatic membrane and adhesive disk, enlarged periplasmic vacuoles and loss of cytoplasmic content (Figure 3 D-F). In some cases, partial internalization of the flagella was observed. In the trophozoites surviving 24 h of culture (Figure 3 H-L), a smaller proportion of trophozoites in the treated culture were morphologically altered as previously described while others displayed a morphology similar to those of the control group (Figure 3 G), although some distinctive differences were observed, mainly consisting in an altered vacuole system and swollen adhesive disk.

**Fig. 3.**
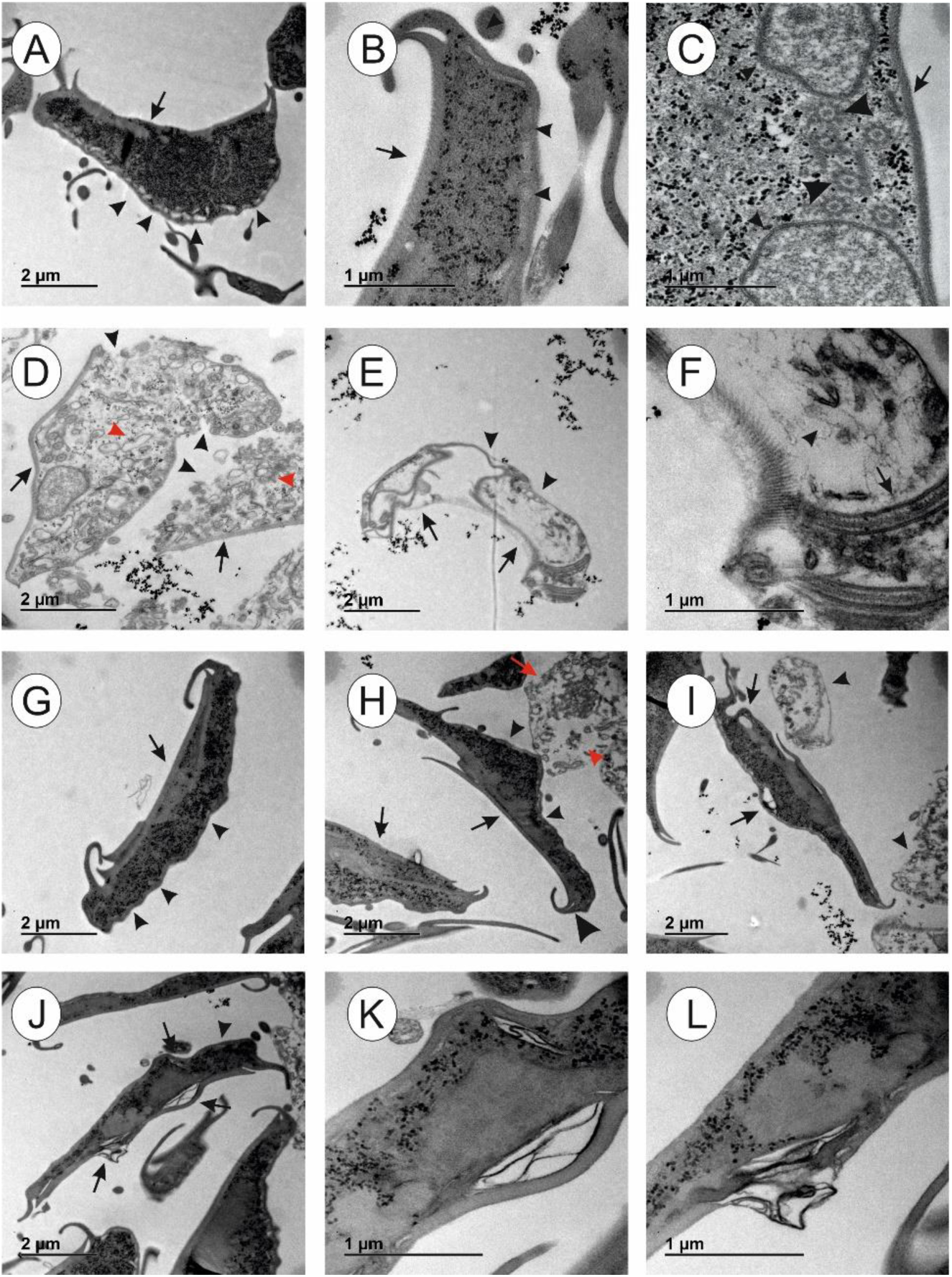
Transmission electron microscopy images of *G. duodenalis* trophozoites from non-exposed group after 1 h culture (A-C) or 24 h culture (G) and γ-terpinene-exposed cultures for 1 h (D-F) and 24 h (H-L). A) Non-exposed trophozoite showing the adhesive disk (arrow) and vesicles under the plasmatic membrane (arrowheads). The cytoplasm has a granular aspect. B) Detail of the adhesive disk (arrow) and the vesicles under the plasmatic membrane (arrowheads). C) Trophozoite with adhesive disk (arrow), flagella with the 9+2 microtubule array (large arrow), and nuclei (small arrowheads). The cytoplasm is granular and the nuclei has a normal morphology. D) Trophozoites after 60 m of exposition to γ-terpinene, showing an apparently normal adhesive disk (arrows), clearly damaged plasmatic membranes (arrowheads), disordered cytoplasm and numerous vacuoles (red arrowheads). E) Trophozoite after 60 m of exposition to γ-terpinene showing alterations in the adhesive disk (arrows) and cytoplasm, partly lost (arrowheads). F) Detail of the cytoplasm in E) showing partial flagella internalization (arrows) and disordered cytoplasm (arrowhead). G) Trophozoite after 24 h of culture, control group, showing normal structures, such as adhesive disk (arrow), vesicles under the plasmatic membrane (arrowheads) and a dense, granular cytoplasm. H) Trophozoites after 24 h of γ-terpinene exposure. Some trophozoites have a normal adhesive disk (black arrow) and vesicles under the cytoplasmic membrane (small black arrowheads), but swollen regions could be observed (large black arrowhead). Altered cells are also present, showing damage in the plasmatic membrane (red arrows) and disordered and diffuse cytoplasm (red arrowheads). I) Trophozoites after 24 h of γ-terpinene exposure. Swollen regions in the cytoplasm and adhesive disk are observed (arrows), as well as clearly altered cells with disordered and diffusing cytoplasm and damage in the plasmatic membrane (arrowheads). J) Trophozoite after 24 h of γ-terpinene exposure, showing normal vesicles under the plasmatic membrane (arrowheads) as well as swollen regions in the adhesive disk and under the plasmatic membrane (arrows). K) and L) show details of the damaged regions shown in J).

## 4. Discussion

Natural products from Lamiaceae and Asteraceae plants are the most widely employed to treat gastrointestinal disorders in traditional medicine across the Mediterranean basin, including diarrhea and infection caused by viruses, bacteria or parasites (Rahman et al., 2016, Redouan et al., 2022). Also, and in a more specific approach, several natural products from plants have been evaluated against *Giardia* spp., with Lamiaceae (30% of the studies) and Asteraceae (13,5%) being the most widely employed (Alnomasy et al., 2021). In our study, several EOs from Lamiaceae displayed excellent AG activity, including Lamiaceae EOs from *Mentha*, *Thymus*, *Lavandula*, *Satureja* and *Origanum* genera.

*M. suaveolens* EOs displayed an excellent AG activity (IC_50_ < 25µg/ml), with results comparable to other *Mentha* species extracts, such as dichloromethane, methanolic and hexane extracts from *M. x piperita*, with no toxic effects (Vidal et al., 2007). However, the composition of the extracts was not described in the above-mentioned study, which made the comparison between studies difficult. *Mentha longifolia* ethanolic extracts, a non-toxic product, also displayed a good IC_50_ value, with menthol as the main component (El-Badry et al., 2010). In our study, piperitenone and piperitenone oxide predominate in the composition of both EOS, but, unfortunately, they could not be assessed against *Giardia* trophozoites.

Excellent AG activity was also obtained with EOs from *T. vulgaris* (IC_50_ < 25 µg/ml) and EOs from *T. zygis* (IC_50_ = 66.9-71.5 µg/ml), and *O. virens* HD EO (IC_50_ = 199.4 µg/ml). Comparable results were obtained by other authors with *Thymus* and *Origanum* species, such as *T. capitata*, *T. zygis silvestris* and *O. virens* (IC_50_ = 71-185 µg/ml) (Machado et al., 2010a), and *Origanum* EOs (IC_50_ = 22-108 µg/ml) (Andrade-Ochoa et al., 2021). In these studies, the main composition of the EOs were based mainly on thymol, γ-terpinene, ρ-cymene, and carvacrol, the most abundant compounds obtained also in the present study in both genera EOs.

In our study, *Satureja* EOs also displayed excellent AG activity with IC_50_ values of <25-109.2 µg/ml depending on the extraction method. Both EOs were rich in carvacrol, thymol, γ-terpinene, and ρ-cymene, compounds that are among the most active against *Giardia* trophozoites. Combinations of carvacrol with γ-terpinene displayed synergistic activity, which could explain the high AG activity detected with the two S*. montana* EOs. A similar composition was found by other authors in *S. montana* EO with anti-bacterial properties (Cavar et al., 2008). In other studies, *Satureja* EOs were found also active against *T. gallinae* (Bailen et al., 2022), *Leishmania* spp. (Monzote et al., 2012), *Acanthamoeba spp., Trypanosoma* spp., fungus and bacteria (Jafari et al., 2016). *Lavandula luisieri* EOs and *L. lanata* HD EO displayed excellent AG activity (IC_50_ = 25-32.1 µg/ml). Two of the *L. intermedia* EOs displayed good AG activity (*L. intermedia* “super” HD EO, IC_50_= 111.3 µg/ml and *L. intermedia* “abrial” HD EO, IC_50_ = 113 µg/ml). There are not many studies in literature evaluating the anti-parasitic activity of *Lavandula* spp. against *Giardia* spp., although previous studies demonstrated a good AG activity employing *L. angustifolia* and *L. intermedia* EOs (Moon et al, 2006). The anti-protozoa activity has been proved also against *Trichomonas* spp. and *Hexamita* spp. (Moon et al., 2006, Bailén et al., 2022). Maybe the composition of the EOs or the synergies between compounds was the key to explaining the differences found. In the composition of *L. luisieri* EOs, trans-α-necrodyl acetate and camphor predominate, while linalool and linalyl acetate predominate in *L. intermedia*, with lower amounts of camphor and 1,8-cineole (Table 3). However, we could not assess trans-α-necrodyl acetate against *Giardia* trophozoites.

*Salvia* species extracts and their components have been previously shown AG activity by other authors (Calzada et al., 2015). In our conditions, *S. sclarea* HD EO displayed good AG activity (IC_50_ = 88.2 µg/ml), with lavandulyl acetate and linalool predominating in the active EO composition, like in the *L. intermedia* EOs of the present study. According to other studies, *Salvia* EOs are active also against other protozoa, such as *Plasmodium* spp. (Monzote et al., 2012).

The main compounds identified in the EOs from plants with antiprotozoal activity are monoterpenoids (including linalool, 4-terpinenol, thymol, carvacrol, citral, limonene, α-pinene, γ-terpinene, α-phellandrene, and *p*-cymene), sesquiterpenes (including β-caryophyllene, nerolidol, α-copaene, cyperene, and germacrene D) and phenylpropanoids (including eugenol, methyl chavicol, and cinnamaldehyde) (Monzote et al 2012). In our study, the most active against *Giardia* trophozoites and less cytotoxic were monoterpenoids, including γ-terpinene, thymol, carvacrol, and α-pinene, as well as the sesquiterpene caryophyllene oxide. All of them displayed a good SI, although α-pinene was slightly cytotoxic for Vero cells. In previous studies employing *T. gallinae*, satisfactory results were also obtained with monoterpenoids (thymol, γ-terpinene, ρ-cymene) (Bailén et al., 2022). Our results regarding terpenes agree with recent studies, in which thymol and carvacrol were active against *Giardia* trophozoites, with IC_50_ values of 21.4 and 31.9 µg/ml (Andrade-Ochoa et al., 2021). Indeed, thymol and carvacrol are isomers with a single change in the position of the hydroxyl group in the phenolic ring. Beta-caryophyllene was one of the active compounds found in the present study and in previous studies conducted by our group with *T. gallinae* (Bailén et al., 2022). This sesquiterpene was also found in extracts from other plants, such as *A. conyzoides* in percentages close to 20%, although other compounds, such as the chromenes precone I & II were more abundant, but AG studies are lacking with these two compounds (Pintong et al., 2020). However, the cytotoxicity of β-caryophyllene in Vero cells found in our study excluded this compound for further studies.

Comparing the chemical structures of the tested terpenes it has been observed that oxidation of the double bound 14,5 of β-caryophyllene to produce caryophyllene oxide reduced cytotoxic effects. Also, the presence and specific position of hydroxyl groups on the structure of the aromatic ring of monoterpenes significantly influence their AG activity, as evidenced by the contrasting effects observed between the active compounds thymol and carvacrol, and the less active ρ-cymene. In any of these cases, Lipinskís rule of five is fulfilled, which makes them compounds that can be orally administered in an easy way.

Unfortunately, compounds that are present in high proportions in the most active EOs evaluated in the present study, such as piperitenone and piperitenone oxide from *Mentha suaveolens* extracts, and trans-α-necrodyl acetate and lavandulol in *Lavandula* species could not be assessed. All of them could be responsible for the AG activity in the above mentioned EOs. Indeed, piperitenone oxide previously showed significant anti-protozoa activity against *Trypanosoma* (Guardo et al., 2017).

As we have shown in the present study, synergistic activities could be acting in some of the employed EOs, especially in those where monoterpenoids are more abundant. We have seen synergistic activity in the combinations of several of them, such as γ-terpinene with carvacrol or thymol, or combination of thymol with ρ-cymene. The synergistic activity was also detected in the combinations of linalyl acetate with linalool, and between linalyl acetate and ρ-cymene or thymol.

None of the experiments using linalool combined with camphor, α-terpineol, or 1,8-cineol displayed synergistic activity, except the above mentioned synergy observed with linalyl acetate and linalool combination. These combinations were present in the HD *L. intermedia* “super and “abrial (IC_50_ = 111.3 and 113 µg/ml, respectively), and HD *S. sclarea* (IC_50_ = 88.2 µg/ml), but all of them have linalyl acetate + linalool in their composition, which could explain the AG activity. Also, no synergistic effect was observed with the combination camphor + 1,8-cineol, present in some of the above-mentioned Lavandulas. Indeed, 1,8-cineol was previously identified acting in an antagonistic manner (Guardo et al., 2017).

The synergistic combinations linalyl acetate + linalool was present in several of the active EOs, *S. sclarea* HD EO (IC_50_ = 88.2 µg/ml), *L. intermedia* “super” HD EO (IC_50_ = 111.3 µg/ml), and *L. intermedia* “abrial” HD EO (IC_50_ = 113 µg/ml).

No synergistic effect was observed with the combination thymol + carvacrol, present in significant amounts in both *S. montana* EOs, although the AG effect could be explained for the presence of the other terpenes, γ-terpinene and ρ-cymene combined with thymol or carvacrol. Indeed, the expected and obtained values of γ-terpinene and ρ-cymene combination reached values close to 100%, making the analysis of synergies with these two compounds difficult.

The most consistent synergies were observed with combinations of the following terpenes, thymol, carvacrol, ρ-cymene, and γ-terpinene. Thymol + γ-terpinene displayed a slight synergistic effect, and they were present in both EOs of *S. montana* (IC_50_ = 25-109.2 µg/ml), *T. zygis* (IC_50_ = 66.9-71.5 µg/ml), and *O. virens* (IC_50_ = 199.4-275.9 µg/ml). The combination thymol + ρ-cymene displayed slight synergism between both compound, and they were present in significant amounts in both EOs of *T. vulgaris* (IC_50_ < 25 µg/ml), *S. montana* (IC_50_ = 25-109.2 µg/ml), and *T. zygis* (IC_50_ = 66.9-71.5 µg/ml). The combination of γ-terpinene and ρ-cymene yielded good AG effects, as we have described above, and they were present at different proportions in both EOs from *T. vulgaris, T. zygis*, and *S. montana*, highly active against *Giardia* trophozoites, and in the less AG active EOs from *O. majorana* (HD EO) and *O. virens* (SD EO). Finally, the combination γ-terpinene and carvacrol displayed good synergistic effects, and they were present at significant amounts in both EOs of *S. montana* (IC_50_ = 25-109.2 µg/ml). There are not many studies of synergies to support our findings, but our results agree with those obtained by other authors who found synergies between carvacrol and γ-terpinene in several EOs, while 1,8-cineole was present in the antagonistic combinations (Guardo et al., 2017). Indeed, the presence of 1,8-cineole was frequently observed in the less AG EOs (Supplementary Table 1). The compounds thymol, carvacrol or γ-terpinene were present at high concentration in the active AG EOs from *Thymus* and *Origanum* species also in other studies (Machado et al., 2010a, Andrade-Ochoa et al., 2021). Besides the positive effect of synergies, these monoterpenes are volatile, lipophilic and difficult to solve, which can help to explain why some EOs had lower IC_50_ than the pure compounds. Terpenoid-rich phenolic extracts appear to target *Giardia* trophozoites through a multi-hit mechanism that combines direct membrane damage with interference in microtubule dynamics (Palomo-Ligas et al., 2022). In addition to their possible direct cytotoxic effects, damage to the ventral disc may impair the parasite’s ability to attach to the intestinal epithelium, thereby promoting its clearance in vivo through intestinal peristalsis.

In our study, γ-terpinene treatment induced significant morphological alterations as early as 60 minutes after exposure, including disruption of the plasma membrane and peripheral vacuoles, deformation of the ventral disc, cytoplasmic disorganization, and flagellar internalization. Similar effects have been described following exposure to methanolic extracts of *Lippia* spp., a plant rich in terpenoids structurally related to γ-terpinene, such as thymol and *p*-cymene (Ponce-Macotela et al., 2006). A similar effect was also observed using extracts from commercial spices containing primarily eucalyptol, cumin aldehyde, and anethole (Andrade-Ochoa et al., 2021). Likewise, monoterpene phenols, mainly thymol and carvacrol, and their hydrocarbon precursors *p*-cymene and γ-terpinene, which are present in aromatic plants such as *Thymbra capitata*, *O. virens*, *T. zygis*, and *Lippia graveolens*, have been shown to permeabilize the trophozoite plasma membrane within minutes, leading to osmotic imbalance and massive vacuolization (Machado et al., 2010a,b; Machado et al., 2011). These membrane lesions have been attributed to the insertion of lipophilic monoterpenes into the lipid bilayer, resulting in increased membrane fluidity and destabilization (Machado et al., 2010a; Andrade-Ochoa et al., 2021).

Other natural products, including curcumin (Pérez-Arriaga et al., 2006), galactoglycerolipids from *Oxalis corniculata* (Manna et al., 2010), garlic extracts (notably allyl alcohol and diallyl thiosulfinate) (Harris et al., 2000; Argüello-García et al., 2018), dichloromethane and methanolic extracts of *Mentha × piperita* (Vidal et al., 2007), and ethanolic crude extracts from *Ageratum conyzoides* (Pintong et al., 2020), have produced similar cytoplasmic and membrane alterations in *Giardia* trophozoites.

Beyond membranolysis, cell damage may result from interactions with tubulin, leading to decreased abundance of α-tubulin and disruption of microtubule organization, particularly within the ventral disc (Palomo-Ligas et al., 2022). Curcumin, for instance, has been shown to bind at the interface of the tubulin dimer, near the colchicine/vinblastine binding site, disrupting cytoskeletal structures including the ventral disc and flagella (Gutiérrez-Gutiérrez et al., 2017).

The ventral disc alterations observed in γ-terpinene-treated trophozoites in the present study are consistent with microtubule destabilization. However, in some studies, no visible damage to cytoskeletal structures or the adhesive disc was observed following treatment with ethanolic extracts of *Lippia* (Ponce-Macotela et al., 2006) or allicin from garlic (Argüello-García et al., 2018), suggesting that cytoskeletal disruption may be compound- or concentration-dependent.

## Conclusions

The EOs of *Lavandula*, *Mentha*, *Thymus,* and *Satureja* genera, plants frequently employed in traditional medicine to treat gastrointestinal disorders, have shown good AG activity and could be used as natural antigiardial agents. The most active compounds identified in the present study, γ-terpinene, thymol, carvacrol and caryophyllene oxide adheres to the Lipinski’s five rules, are non-toxic, and the first three of them have been authorized for use as additives for animals food (Commission Implementing Regulation (EU) 2024/1989 of 22 July 2024), which makes them ideal for alternative or complementary treatment of giardiasis, and probably other gastrointestinal protozoosis.

## Authors contribution

Conceptualization and design: MB, AG and MTG. Acquisition of data: SMH, SAF, MJI, JAD, FP-G, JNR and IAC. Analysis and interpretation of data: MB, AG, MTG, SMH, SAF, MJI, JAD, FP-G, JNR and IAC. Funding acquisition: AG and MTG. Draft of the article: MB and MTG. Critical revision for important intellectual content: MB, AG and MTG. Final approval: all the authors.

## Abreviations

AG: Anti-*Giardia*
ATCC: American Type Culture Collection
CC_50_: Cytotoxic Concentration 50
CITA: Centro de Investigación y Tecnología Agroalimentaria de Aragón
DMSO: Dimethyl sulfoxide
ECDC: European Center for Disease Control
EO: Essential Oil.
EOs: Essential Oils
EPA: Environmental protection Agency
EU: European Union
GC: Gas Chromatography
GC-MS: Gas Chromatography-Mass spectrum
HD: Hydrodistillation
HD EO: Essential Oil obtained by hydrodistillation
IC_50_: Inhibitory concentration 50
ICA-CSIC: Instituto de Ciencias Agrarias, Centro Superior de Investigaciones Científicas
IPE-CSIC: Instituto Pirenáico de Ecología, Centro Superior de Investigaciones Científicas (Spain)
MTT: 3-(4,5-Dimethylthiazol-2-yl)-2,5-Diphenyltetrazolium Bromide
NIH: National Institutes of Health
NIST: National Institute of Standards and Technology
NIST 17: NIST/EPA/NIH Mass Spectral Library
PMS: Phenazine methosulfate
SD: Steam distillation
SD EO: Essential Oil obtained by Steam Distillation
SI: Selectivity index
TEM: Transmission Electron Microscopy
TYI-S-33: Tripticase-Yeast extract-Iron modified by Diamond and others

## Supplementary material

**Supplementary Figure 1.**
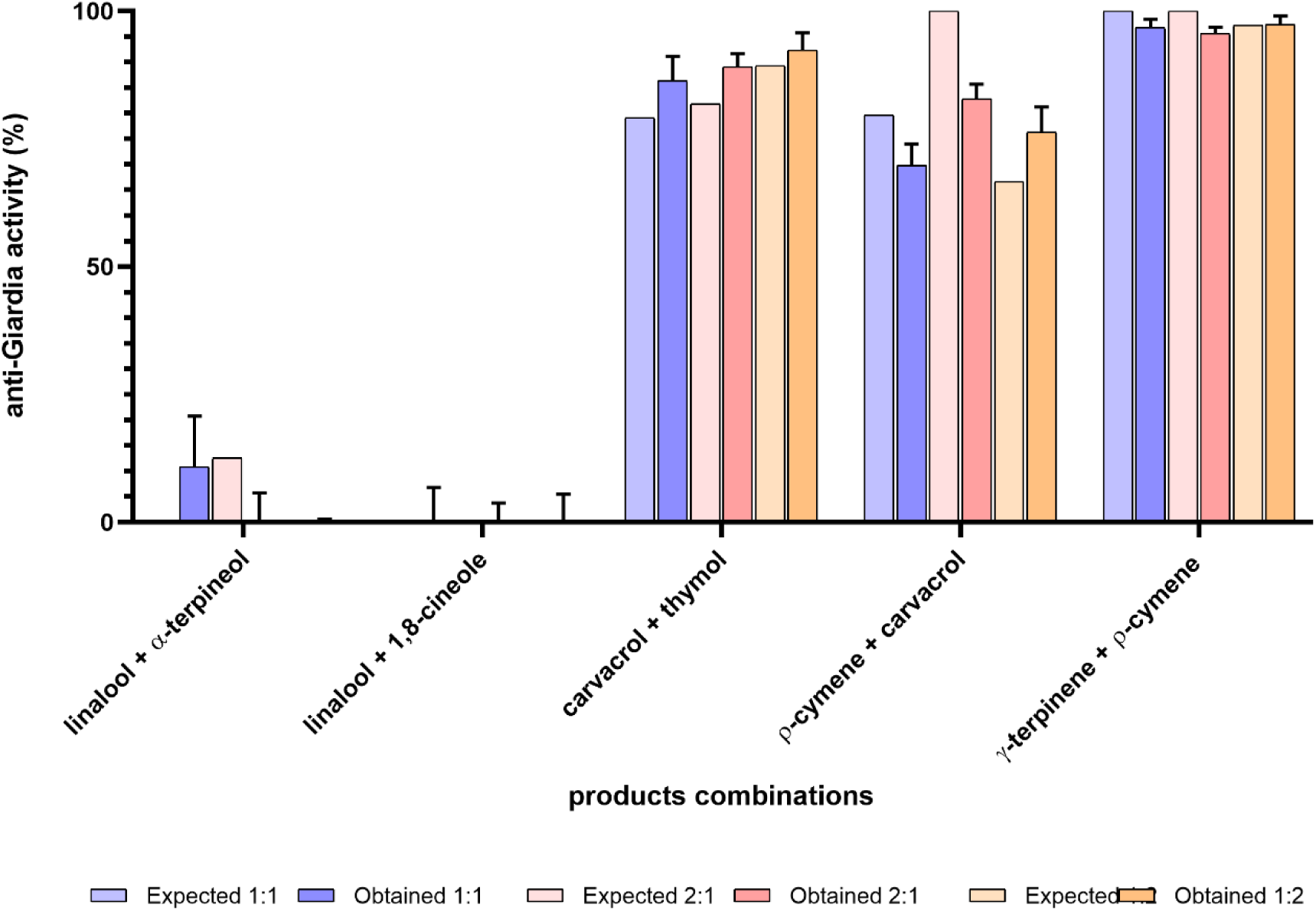
Activity of compounds combined in pairs from the most active AG EOs that did not show synergistic activity.

**Supplementary Figure 2.**
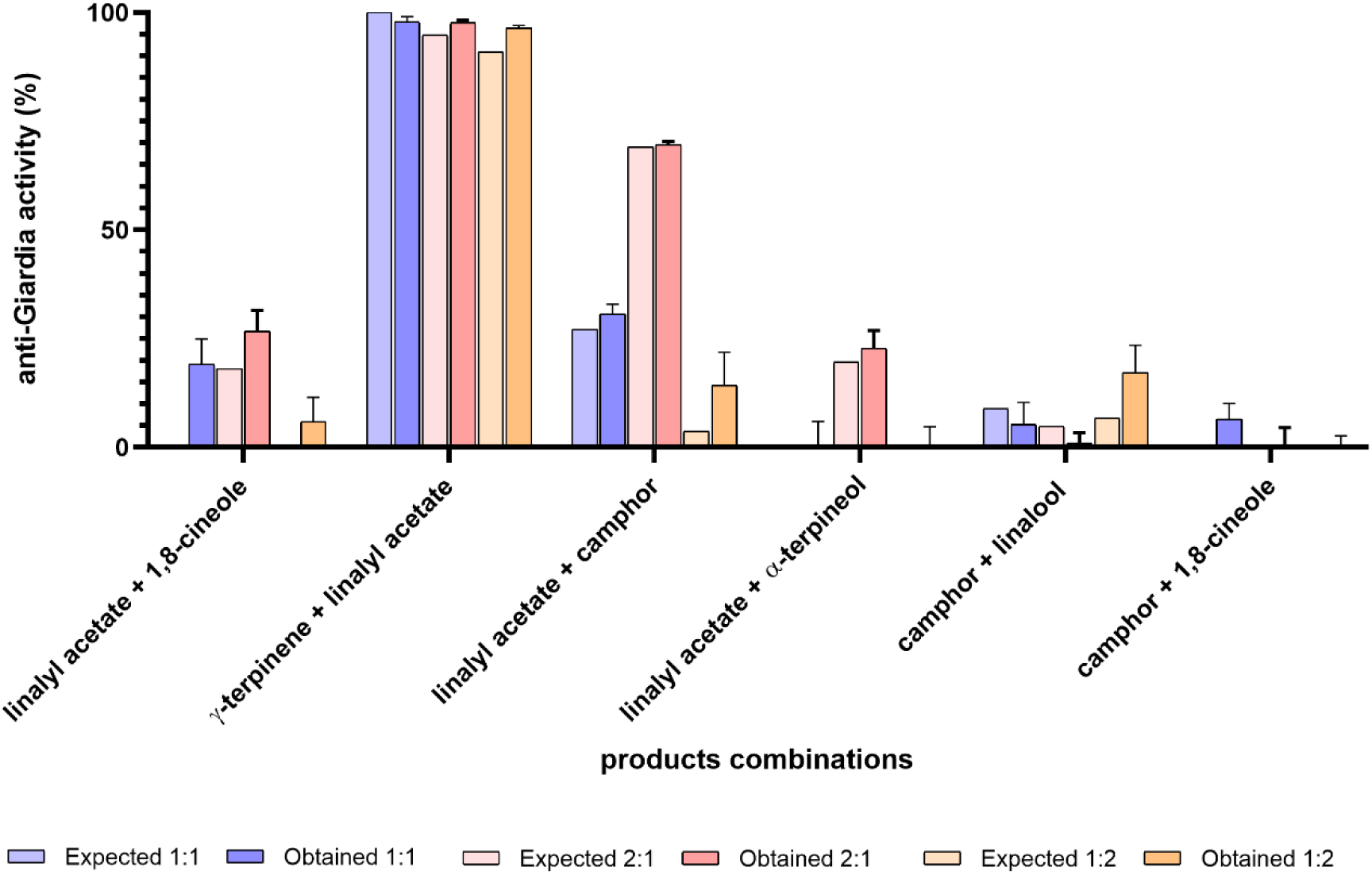
Activity of compounds combined in pairs from the most active AG EOs that did not show synergistic activity.

**Supplementary Table 1.**
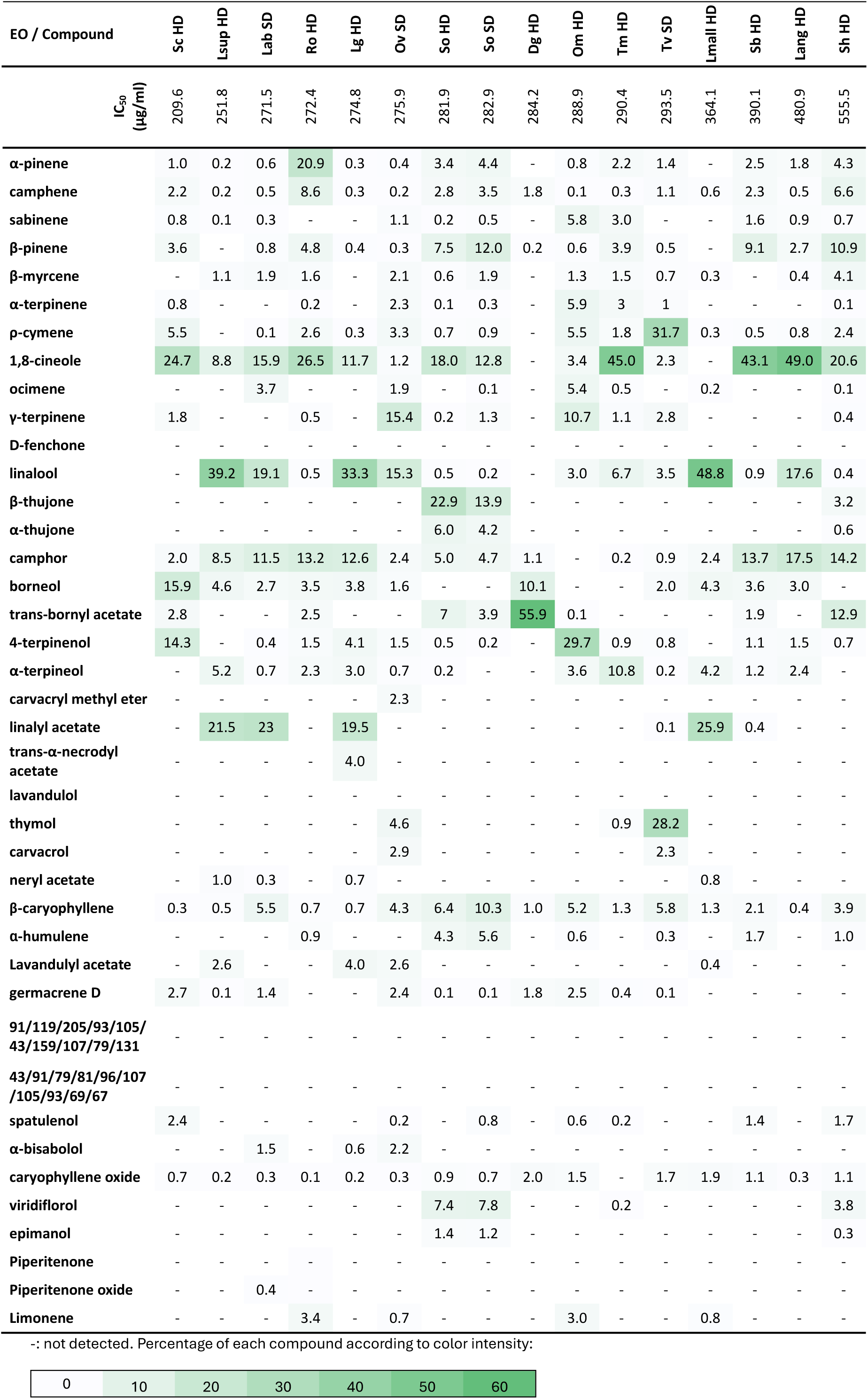
Main components (in percentage) of the EOs with IC_50_ values higher than 200 µg/ml and ordered by retention index. Sc: *S. chamaecyparissus*; Lsup: *L. intermedia* “Super”; Lab: *L. intermedia* “Abrial”; Ro: *R. officinalis*; Lg: *L. intermedia* “Grosso” ; Ov: *O. virens;* So: *S. officinalis*; Dg. *D*. *graveolens*; Om: *O. majorana*; Tm: *T. mastichina*; Tv: *T. vulgaris*; Lmall: *L. angustifolia var. mallete*; Sb: *S. blancoana*; Lang: *L. angustifolia*; Sh: *S. hybrid*. HD, EOs obtained by hydrodistillation; SD, EOs obtained by steam distillation.

